# The long-term sand crab study: phenology, geographic size variation, and a rare new colour morph in *Lepidopa benedicti* (Decapoda: Albuneidae)

**DOI:** 10.1101/041376

**Authors:** Zen Faulkes

## Abstract

Little is known about the basic ecology of albuneid sand crabs because they conceal themselves by digging in sand, and are typically found at low densities. I sampled sand crabs, *Lepidopa benedicti* were sampled at South Padre Island, Texas, regularly for more than five years. Density is high in the summer, and low in the winter. The sex ratio is slightly, but consistently, female skewed. Ovigerous females, carrying about one thousand eggs, are found in mid-summer, with most of the young of the year settling in autumn. The average size of individuals in the South Texas population is smaller than individuals in the Atlantic Ocean, but the population density appear to be higher in Texas. I also describe a new orange colour morph. The results are not consistent with an earlier suggestion that South Texas acts as a population sink for *L. benedicti*.

## INTRODUCTION

Sand crabs (Family Albuneidae) are small decapod crustaceans that live in fine sandy beaches (Boyko 2002). All species in this family are obligate diggers as adults. They conceal themselves completely in sand (Faulkes & Paul 1997; Faulkes & Paul 1998; Dugan, Hubbard & Lastra 2000), and leave no traces of their presence visible to observers walking on a beach. Sand crabs can be difficult to find, with Hay & Shore (1918) noting that they could not find a live *Lepidopa websteri* individual even after “a vast amount of digging”

*Lepidopa benedicti* is a representative albuneid species that lives in the Gulf of Mexico and the Atlantic coast of Florida (Fig. 1). As part of an ongoing long term study to understand the basic biology of a representative species, I have been studying the biology of the species on South Padre Island, Texas (Nasir & Faulkes 2011; Murph & Faulkes 2013; Joseph & Faulkes 2014). This paper more than doubles the time frame of this long term study (from two years to five years), thus providing greater resolution into the phenology of this sand crab, and more opportunities to record rare events. Additionally, this paper provides data on *L. benedicti* from the Atlantic coast of Florida as an initial effort to characterize differences in this population across a wide portion of its range.

**Figure 1:**
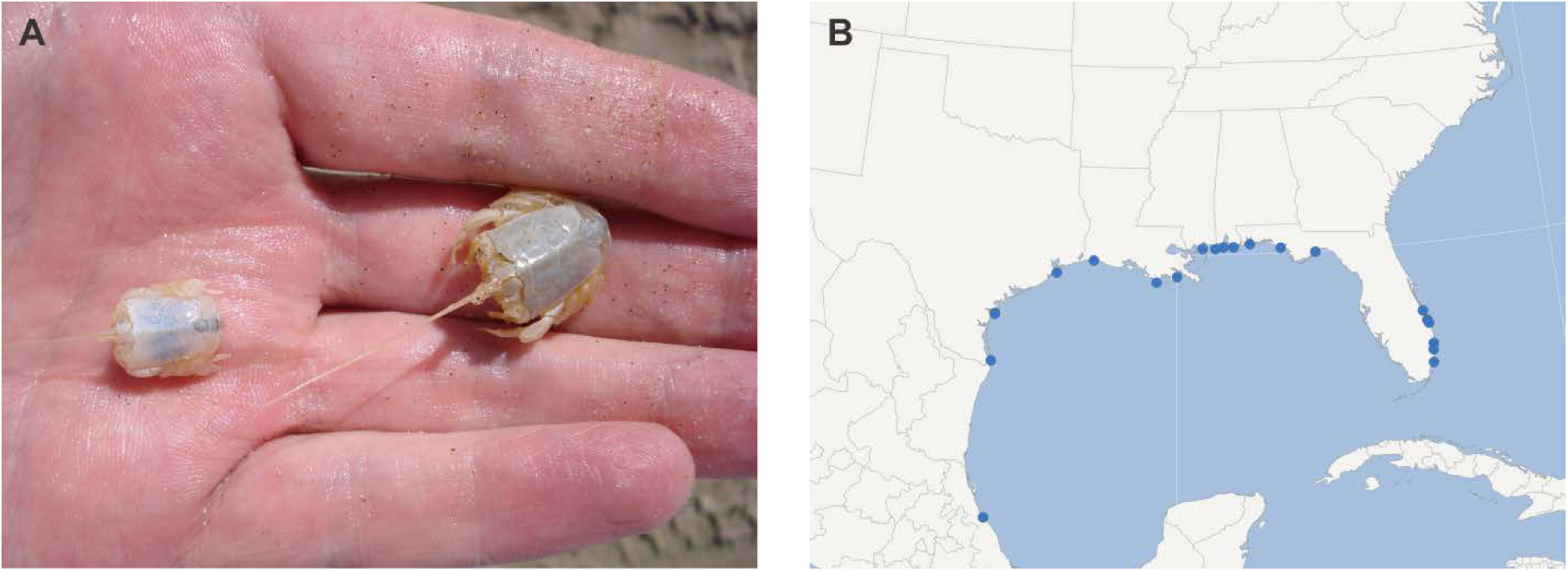
Lepidopa benedicti. (A) Two *Lepidopa benedicti* individuals, showing colour variation. Anterior is to the left. (B) Distribution of *L. benedicti*, based on data from (Boyko 2002); one site shown for each U.S. county or parish.

Data from the first paper from this project (Murph & Faulkes 2013) prompted questions that can be answered with a larger data set. (Murph & Faulkes 2013) suggested that South Padre Island, Texas might be a population sink for sand crabs, for two reasons. First, no ovigerous females had been found over two years of sampling. Continued sampling allowed me to test the possibility that ovigerous females were simply uncommon. Second, individuals from South Padre Island were smaller than other areas where the species is distributed, according to archival records in (Boyko 2002). However, it is possible the ovigerous because the data from other locations were collected by many individuals over many decades, it was possible that such size differences reflected researchers’ preservation biases rather than being a representative sample of the population. Sampling from the Atlantic coast of Florida, and recording all individuals there, could distinguish between preservation bias and a biological difference in size.

Portions of this work have appeared in brief (Faulkes & Feria 2012; Real Scientists 2014).

## METHODS

Sand crabs, *Lepidopa benedicti* Schmitt, 1935, were collected at South Padre Island, Texas, monthly from to August 2009 to December 2014 (exception: data from July 2010 lost). The first two years of data are available at (Faulkes 2014) and were analyzed in (Murph & Faulkes 2013). *Lepidopa benedicti* was also collected at Fort Lauderdale, Oakland Park, and Pompano Beach in southern Florida in November 2012, using the same methods as the Texas site. Results refer to the Texas population unless otherwise specified.

I collected crabs by digging 10 m transects parallel to, and slightly above, the waterline of the swash zone. Three or more transects were dug every month. I overturned sand and examined it for *L. benedicti*, and also collected any individuals which emerged from the trench when it filled with water. I recorded the sex, carapace length, and colour on site. Sample sizes for these features vary because some sand crabs were damaged by the shovel during collecting, escaped, and so on. For example, some damaged animals could be sexed but not have the carapace length measured.

Carapace lengths for *L. websteri*, *Albunea gibbesii*, and *A. catherinae* were taken from (Boyko 2002).

Graphs and statistical analyses were done using Origin 7 SR2 (OriginLab Corporation, Northampton, Massachusetts). Graphs showing values per month show average of all transects per month per year, to avoid pseudoreplication. That is, sample size for most months is five, one for each year of the project.

## RESULTS

### Abundance

The mode abundance of *L. benedicti* per transect is less than 1 10 m^−1^ transect (Fig. 2A). The abundance of *L. benedicti* varies throughout the year, however, being low in winter and peaking in summer (Fig. 2B). This confirms and refines previous results (Murph & Faulkes 2013).

**Figure 2:**
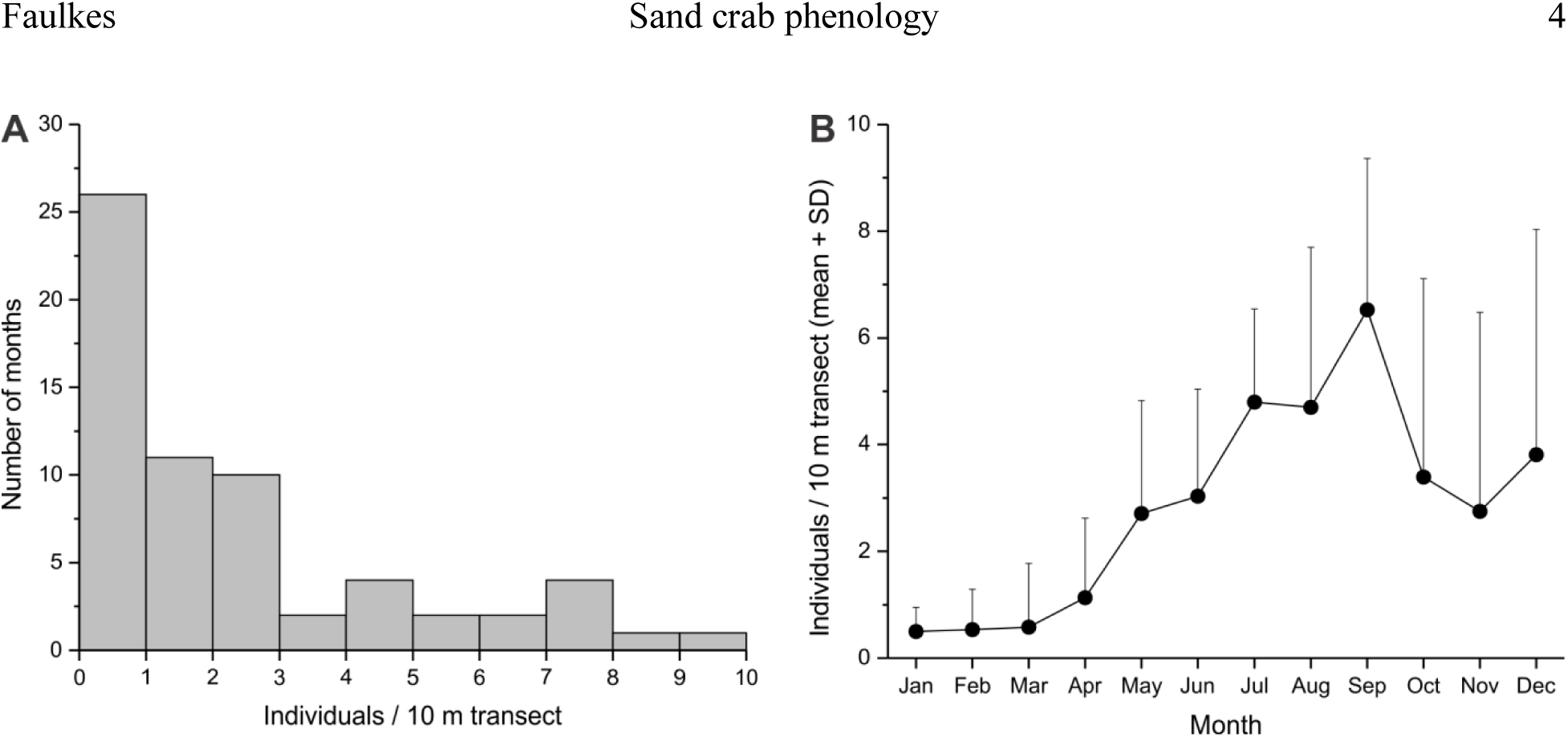
Lepidopa benedicti abundance. (A) Histogram of abundance of *Lepidopa benedicti* per 10 m transect of beach. (B) Abundance of *L. benedicti* per month.

The abundance of individuals in Florida was less than South Padre Island, Texas, although this difference could not be analyzed statistically. In November, the abundance of *L. benedicti* was 2.75 individuals 10 m^−1^ transect on average (S.D. = 3.73, n = 6) in Texas, but was 0.71 individuals 10 m^−1^ transect in Florida (n = 1), even with fewer transects dug in Texas overall (36 transects) than Florida (42 transects).

### Size

The average carapace length of individuals was 8.44 mm (SD = 2.14), with the largest individual measuring 17.92 mm, and a minimum of 2.42 mm. The size distribution is unimodal. Females are slightly, but significantly, larger than males (Fig. 3A; t_752_= −6.97, p < 0.01), confirming previous results (Murph & Faulkes 2013).

**Figure 3:**
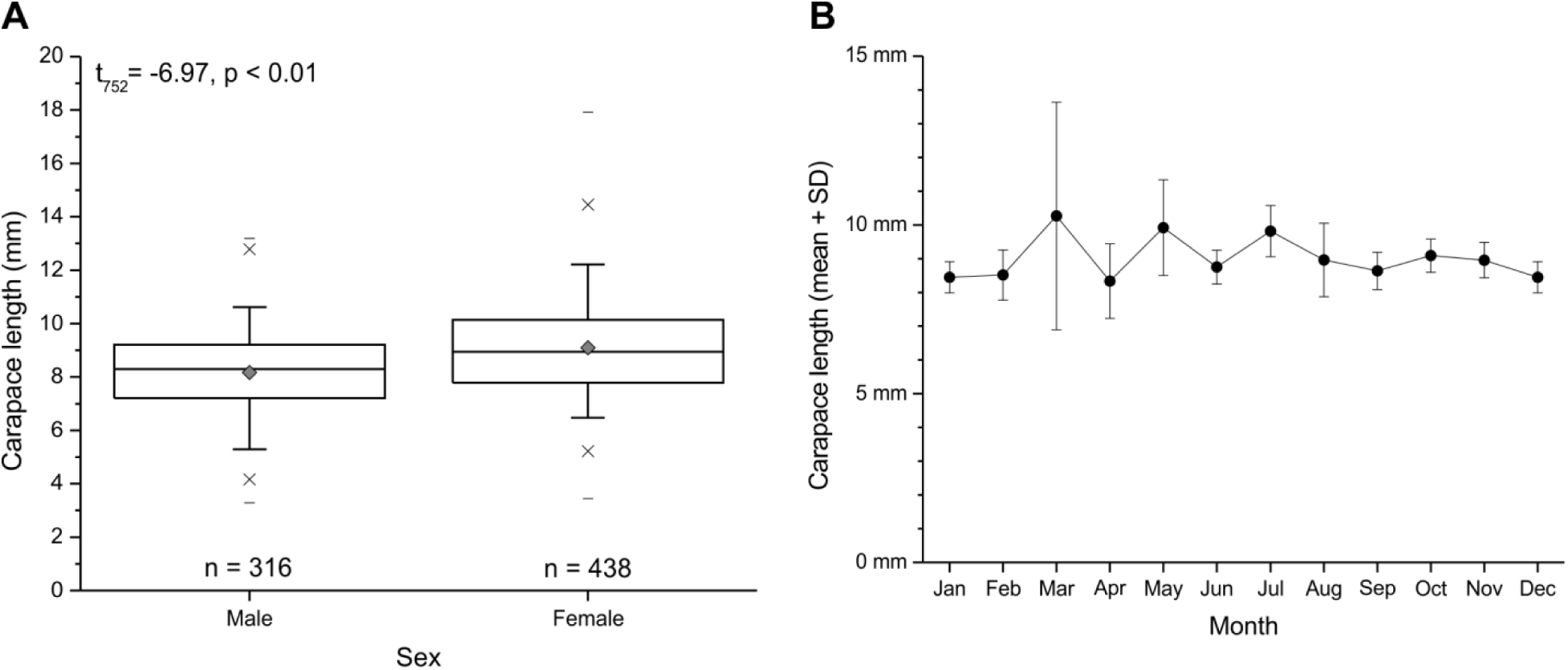
Size of Lepidopa benedicti. (A) Box plot of size of both sexes. (B) Size of individuals per month excluding young of the year (i.e., those less than 5 mm carapace length).

One possible explanation for the variation in abundance is that adult mortality and/or recruitment of young (below) is strongly seasonal. One prediction of this hypothesis would be that there would be annual variation in size, with the average size dropping if the largest (and presumably oldest) individuals die seasonally. Annual changes in size of adults were examined by averaging the carapace length of individuals excluding those one standard deviation below the mean (i.e., assumed to be sub-adult individuals). The mean size of adult individuals does not vary over the year (Fig. 3B).

*Lepidopa benedicti* collected from Florida were not significantly larger than Texas individuals (Fig, 4A; t_11_=1.99, p=0.072), controlling for sex (all female) and month (all collected in November).

**Figure 4:**
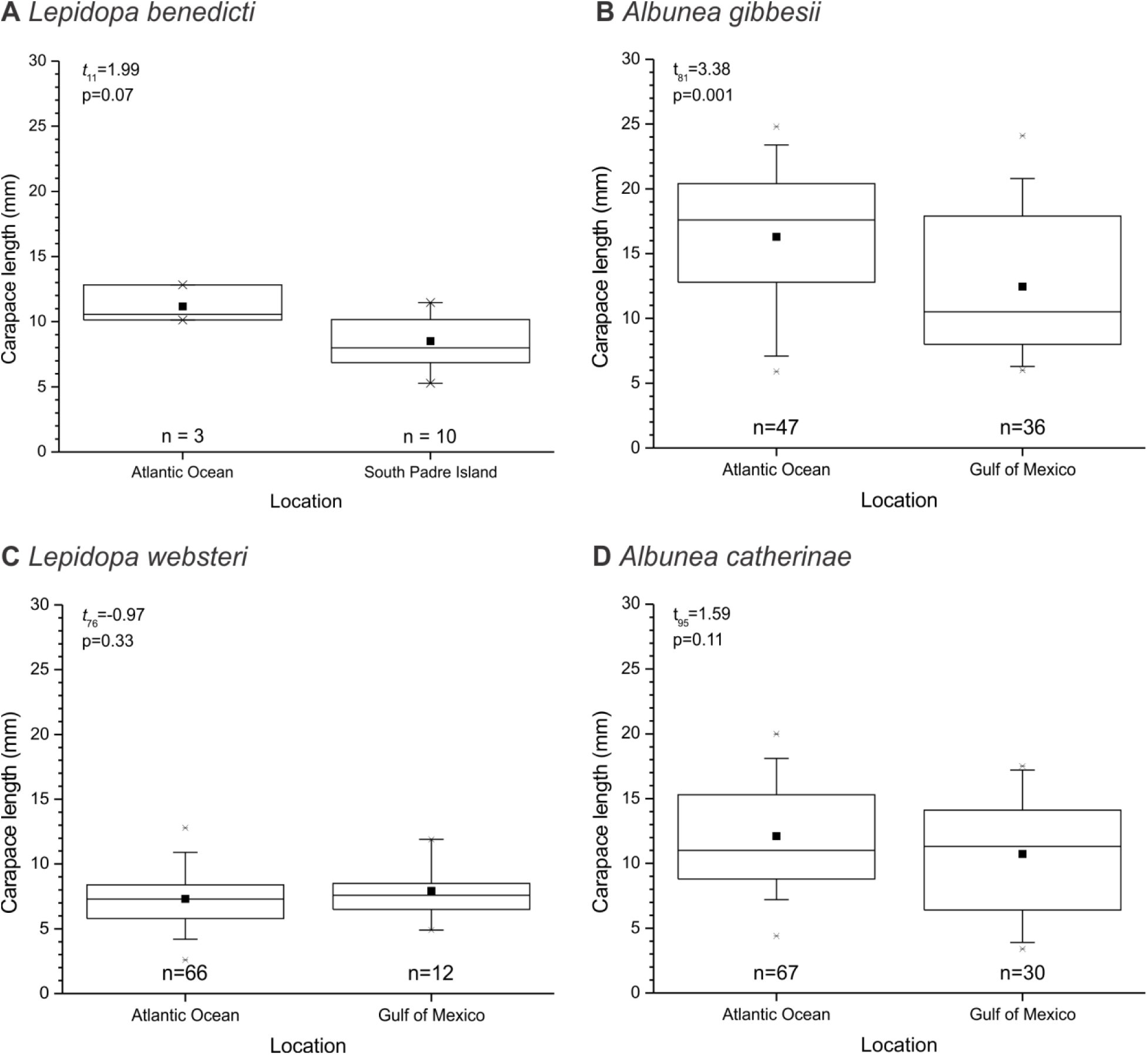
Comparison of sand crab sizes from Gulf of Mexico and Atlantic Ocean. Size of sand crabs collected from Atlantic Florida and Gulf of Mexico. (A) *L. benedicti*, this study, matching sex and month. (B) *Lepidopa websteri*. (C) *Abunea gibbesii* (D) *Albunea catherinae*. Data for B-D from (Boyko 2002).

All albuneid species are obligate diggers, and presumably have generally similar basic biology. If the western Gulf of Mexico is generally a poor habitat for sand crabs relative to the Atlantic Ocean, other albuneid species might show the same differences in size between the two populations. This was not the case. Like *L. benedicti*, *Albunea gibbesii* collected from the Atlantic Ocean were significantly larger (Fig. 4B; t_81_=3.38, p = 0.001) than those collected from the Gulf of Mexico (Fig. 4B), although the difference was not as pronounced as in *L. benedicti*. There was no significant size difference between individuals collected in the Atlantic Ocean and the Gulf of Mexico in either *Lepidopa websteri* (Fig. 4C; t_76_=-0.97, p=0.33) or *Albunea catherinae* (Fig. 4D; t_95_=1.59, p=0.11).

### Sex and reproduction

The overall sex ratio of *L. benedicti* is skewed towards females, as previously reported (Murph & Faulkes 2013): 58.20% of the crabs were female. The sex ratio varies over the year, however, with males outnumbering females in November and December (Fig. 5).

**Figure 5:**
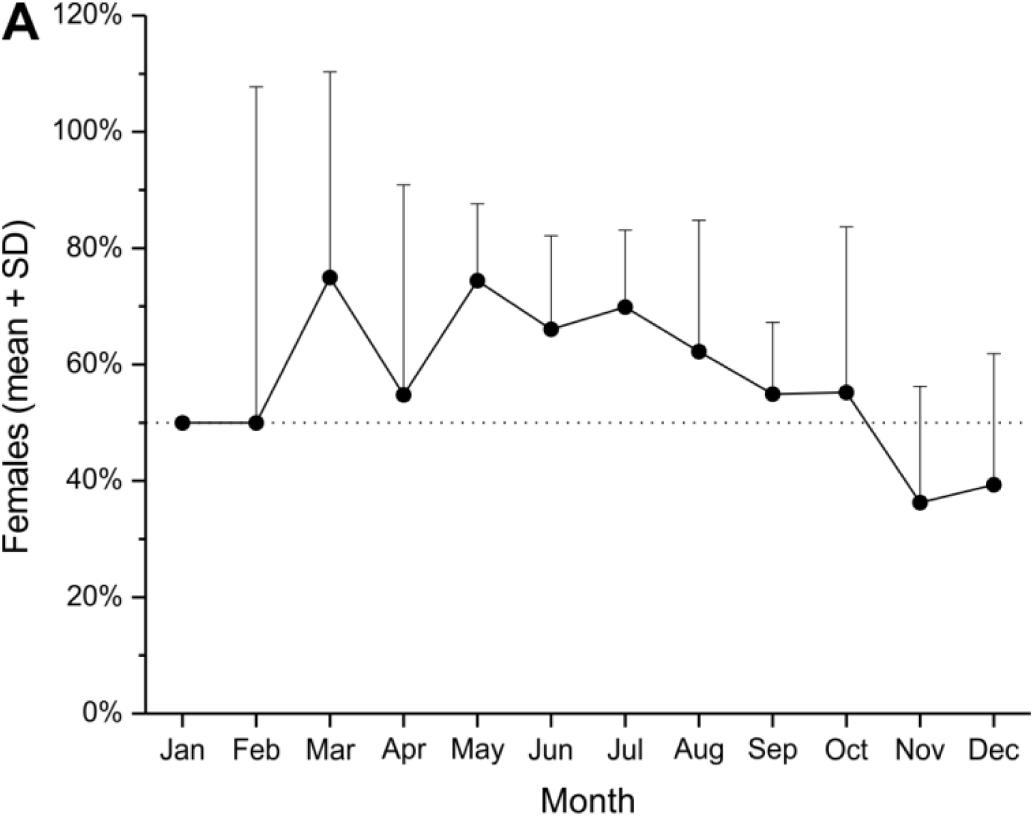
Sex ratio of Lepidopa benedicti. Sex ratio per month.

Three ovigerous females (Fig. 6A) were found in June and August, the same months as other ovigerous *L. benedicti* recorded in the literature (Stuck & Truesdale 1986; Boyko 2002). The earliest was collected on 23 June and the latest on 14 August. I counted the eggs of two individuals. One (9.67 mm carapace length) had 1,419 eggs, and the second (9.16 mm carapace length) had 963 eggs. The latter is close to the smallest ovigerous individual recorded (9.1 mm carapace length; Boyko 2002).

**Figure 6:**
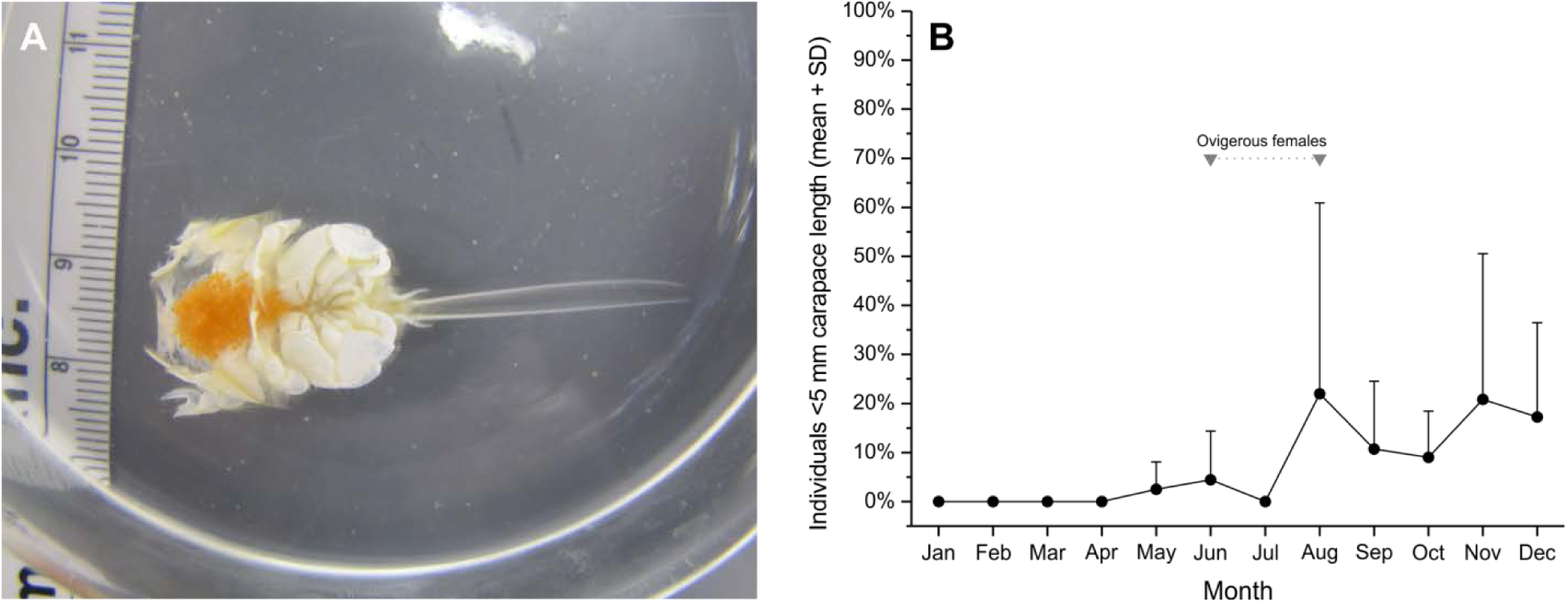
Reproductive aspects of Lepidopa benedicti. (A) Ovigerous female. (B) Young of the year by month.

It is not known how long females retain eggs, but the pelagic larval stages of *L. benedicti* last 14-15 days before individuals metamorphose into benthic megalopae (Stuck & Truesdale 1986). Thus, the young of the year (defined as individuals with a carapace length less than 5 mm) are expected to be found in early autumn. Small individuals are most often found from August to December, and are not found from January to April (Fig. 6B).

### Orange colour morph

Most *Lepidopa benedicti* have a gray carapace, although some individuals are white (Nasir & Faulkes 2011). During the course of this study, I found two orange individuals (Fig. 7). One was a female with an 11.14 mm carapace length collected on 22 December 2013, and the other was a male with a 9.92 mm carapace length on collected on 31 July 2014. The orange colour was visible over the entire dorsal surface, including the ocular peduncles, in both individuals. The pigmented eye spots were the normal black (Fig. 7C) The ventral surface was not noticeably coloured (Fig. 7D), similar to individuals with grey and white carapace (Nasir & Faulkes 2011). The exoskeleton was firm and gave no indication that this animal had recently molted, which can sometimes affect carapace colour. The individual collected in December 2013 remained the same conspicuous orange colour until the animal died on 10 January 2014 from unknown causes.

**Figure 7:**
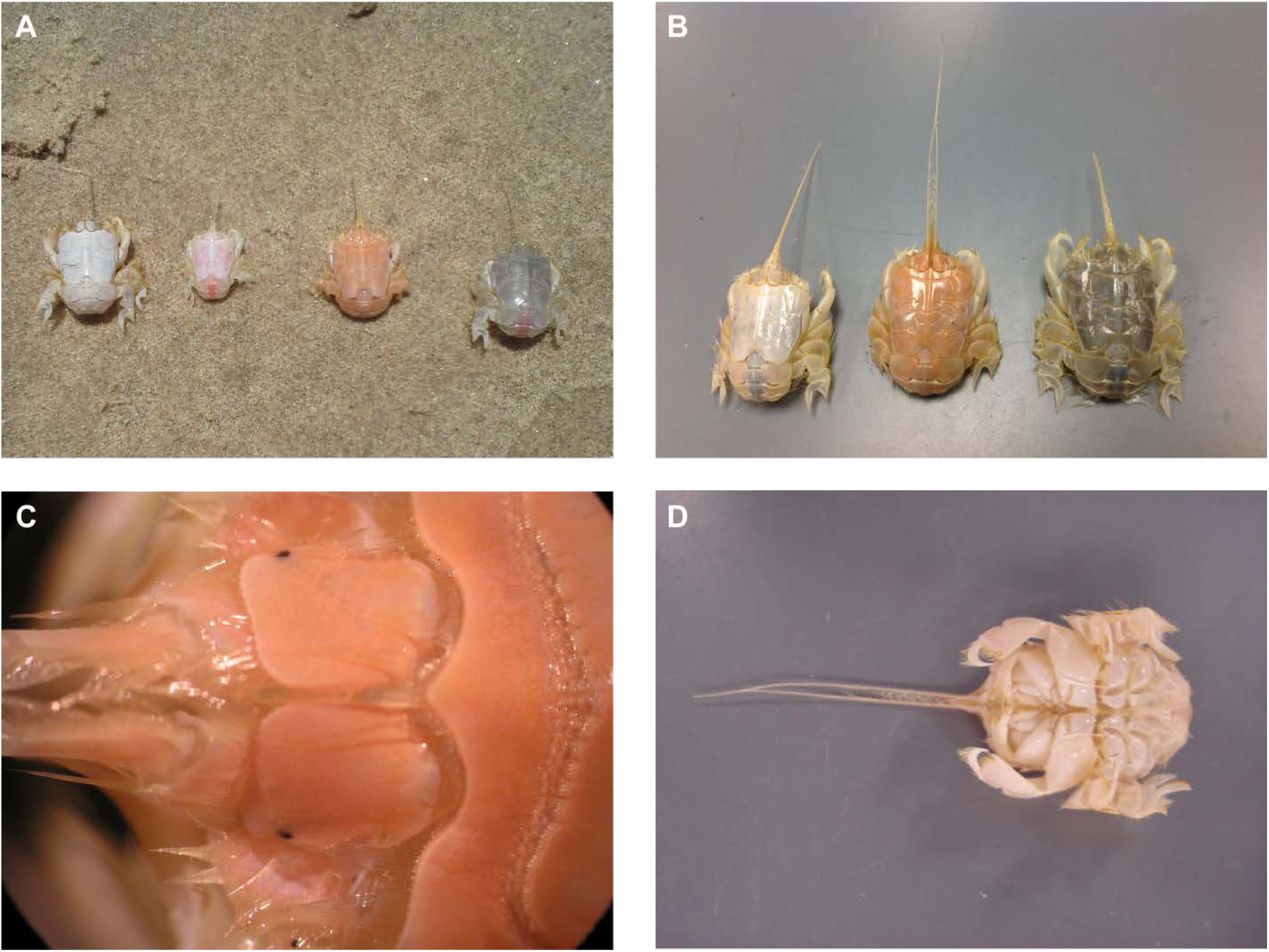
Orange colour morph of *Lepidopa benedicti*. (A, B) Orange individuals next to more common gray and white morphs. Individual in B is same individual shown in C, D, and different than A. (C) Close-up of ocular peduncles, showing typical eyespot pigmentation. (D) Ventral view of orange individual, showing typical white colour of exoskeleton (compare to Fig. 1C in Nasir & Faulkes 2011).

## DISCUSSION

*Lepidopa benedicti* show several annual cycles, despite that this species lives in a subtropical climate with relatively modest weather changes. The available evidence indicates that *Lepidopa benedicti* breed in mid-summer, with recruitment occurring in late summer and autumn. This annual reproductive cycle does not explain the annual changes in abundance, however. Recruitment of the young of the year occurs before abundance drops in January, and there is no evidence of seasonal die off of large individuals at any point in the year. These facts suggest that *L. benedicti* relocate themselves in slightly deeper water below the tide line during the winter months. *Lepidopa benedicti* has been collected in waters up to 3 m deep (Boyko 2002), indicating that this species is not confined to the swash zone.

The hypothesis that South Padre Island is a population sink for *L. benedicti* is not supported by new observations here. The first line of evidence against the sink hypothesis is that *L. benedicti* appear to be more abundant at South Padre Island than in Florida. More sampling of the Atlantic population would be extremely useful, however. Second, ovigerous females are present at South Padre Island, although they are difficult to collect, perhaps because eggs are not retained long before being released. The only recorded hatching of *L. benedicti* eggs occurred within the day of collection (Stuck & Truesdale 1986). Female *L. benedicti* generate about 1,000 eggs (this study). Under lab conditions, 31.25% of larvae survive to the megalopa stage (Stuck & Truesdale 1986), which suggests that about 300 offspring per female could survive to the settlement stage. There is undoubtedly predation in the pelagic portion of the life cycle (which lasts 14-17 days; Stuck & Truesdale 1986) that further reduces larval survival. Continuing this long term study should provide the additional data needed to estimate the population ecology of the species.

The appearance of an orange colour morph in *L. benedicti* is reminiscent of some rare colour morphs of commercially fished crustaceans, such as American clawed lobsters (*Homarus americanus*), which will often make national news (CBC News 2013; Harish 2013; White 2013). In the news media, the probability of finding a red or orange lobster in the wild is usually estimated as being one in 10 million (Lobster Institute 2011; CBC News 2013; Harish 2013; White 2013), although how this estimate has been calculated is unclear. The odds of *L. benedicti* being orange appear to be about 1 in 500. Crustacean colour is determined both by genetics and environment (Kent 1901; Bowman 1942; Black & Huner 1980; Tlusty & Hyland 2005). Some crustaceans can change their colours to some degree (Barnard et al. 2012; Wade et al. 2012), including hippid sand crabs (Wenner 1972; Bauchau & Passelecq-Gérìn 1987), which can change their color to match the sand they live in. The rarity of the orange morph, plus its stability in adults, suggests that this colour is a rare recessive allele or mutation in both *L. benedicti* and lobsters. Because *L. benedicti* are obligate diggers that spend their effectively all their adult lives submerged in sand, colours are unlikely to have any major signalling functions, either to conspecifics, or other species (e.g., predators). Thus, unusual colour morphs in *L. benedicti* may be under less selection pressure than unusual morphs in benthic crustacean, and thus more common. This is consistent with the discovery of this orange sand crab after sampling hundreds of individuals, rather than the millions that might be expected for lobsters.

## ACKNOWLEDGEMENTS

I thank Kevin Faulkes for collecting assistance and Karren Faulkes for emergency shovel replacement.

